# PoolParty: streamlined design of DNA sequence libraries in Python

**DOI:** 10.64898/2026.04.06.716802

**Authors:** Zhihan Liu, Aidan Cordero, Justin B. Kinney

**Affiliations:** Simons Center for Quantitative Biology, Cold Spring Harbor Laboratory, 1 Bungtown Rd., Cold Spring Harbor, New York 11724, United States

## Abstract

**Background:** Computationally designed DNA sequence libraries are essential components of massively parallel reporter assays (MPRAs), deep mutational scanning (DMS) experiments, and other multiplex assays of variant effect (MAVEs). They are also increasingly used in silico to analyze genomic AI models. Designing these libraries, however, remains tedious and error-prone due to the scarcity of purpose-built software.

**Results:** Here we describe PoolParty, a Python package that streamlines the design of complex oligo pools using a simple but flexible API. In PoolParty, each library is represented by a computational graph that can be specified in just a few lines of code. Over 50 built-in operations cover nucleotide- and codon-level mutagenesis, motif insertion, barcode generation, and more. PoolParty automatically generates informative names for each sequence and provides “design cards” detailing how each sequence was generated. Visualization methods let users quickly audit library content and inspect the underlying graph. PoolParty thus transforms oligo pool design from a tedious task requiring custom functions and scripts into a structured, transparent, and reproducible process.

**Conclusions:** PoolParty streamlines the design of DMS, MPRA, and other multiplex assay libraries, and the design cards it provides can help researchers systematically probe and interpret genomic AI models. PoolParty can also be extended to support new assays and analysis strategies as they emerge.

## Background

Designed libraries of DNA sequences (oligonucleotide pools) have become essential tools in functional genomics. In massively parallel reporter assays (MPRAs), complex libraries of DNA sequences are routinely used to measure the regulatory activities of tens of thousands of candidate regulatory elements or their variants in a single experiment [1–13]. In deep mutational scanning (DMS) experiments, coding sequences that have been either systematically or randomly mutagenized are commonly used to map protein fitness landscapes [14–20]. Oligo pools can also be used in high-throughput biochemical assays, such as measurements of transcription factor specificity [21–27]. These experiments are all examples of multiplex assays of variant effect (MAVEs), a large and expanding family of high-throughput methods that quantitatively link sequence to function at scale [28–30]. Beyond the wet lab, designed sequence libraries are increasingly used in silico to probe the behavior of genomic AI models [31–35] and to augment genomic AI training data [36].

Computationally designing these libraries is straightforward in principle, but in practice it can be tedious and error-prone. Typical applications require oligo pools comprising multiple types of variant sequences. A DMS library, for example, might combine exhaustive single-amino-acid substitutions, sampled higher-order mutants, multiple copies of wild-type and control sequences, and barcodes—each generated by a different procedure and subject to different coverage requirements. An MPRA library might tile multiple transcription factor binding sites across a regulatory element in all possible permutations and orientations, then assign each variant multiple barcodes. Coordinating this kind of combinatorial logic in a one-off script quickly becomes unwieldy, and often produces errors that are hard to catch.

Existing tools can help with parts of this process. General-purpose sequence toolkits [37–39] can manipulate sequences but have no notion of a variant library as a structured object. Consequently, users must write custom scripts to handle the combinatorial logic themselves. Several other tools automate library design or sequence perturbation for particular applications. For example, VaLiAnT [40] generates mutational scanning libraries of both coding and noncoding regions; supported mutation types include single-nucleotide substitutions, amino acid substitutions, and in-frame deletions. For MPRAs, MPRAnator [41] provides modules for motif placement, SNP design, and negative control generation [see also 42]. For in silico sequence experiments, tangermeme [43] supports sequence perturbation and analysis of genomic deep learning models. However, each of these tools is targeted to specific applications and, as a result, no individual tool can generate the full range of library designs in common use. For example, VaLiAnT does not support regulatory sequence libraries containing random combinations of transcription factor binding sites, while MPRAnator and tangermeme do not support codon-aware mutagenesis of protein-coding sequences.

Here we present PoolParty, a Python package that enables the streamlined design of complex DNA sequence libraries. In PoolParty, libraries are represented as objects called *Pools*, while individual steps in the sequence generation process are called *Operations*. Using a concise and transparent syntax, users chain together Operations into a directed acyclic graph (DAG)—an approach inspired by deep learning frameworks—that yields the desired Pool. The DAG describes the high-level logic used to generate sequences, while the procedural details and bookkeeping are handled internally. Importantly, no sequences are generated until the user requests them, and it is simple to generate small test sets of sequences. This allows users to explore multiple design options without having to generate complete libraries at each iteration. When sequences are generated, each is automatically assigned an informative name and paired with a *design card* that records the specific procedures used to construct it. We demonstrate PoolParty through three example applications: a DMS library for protein GB1, an MPRA library probing transcriptional regulatory grammar, and an in silico experiment characterizing the genomic deep learning model SpliceAI [44], in which design cards provide the covariates for surrogate modeling.

## Implementation

### Pools and Operations

PoolParty is built around two core abstractions: Pools and Operations. Pools represent collections of sequences—either the final library that the user wishes to generate, or intermediate sets of sequences out of which the final library is constructed. Operations represent the steps used to create Pools. When designing a library, users chain together multiple Operations into a DAG that yields the desired Pool, called the *root Pool*. Sequences are then generated on demand from the root Pool. By delaying sequence generation until it is explicitly requested, users can test many different library designs before committing to generating the full library. For example, users can inspect a few sampled sequences from each one of multiple proposed designs.

PoolParty provides over 50 built-in Operations that users can choose from when constructing DAGs. Each Operation takes zero or more Pools as input and returns exactly one Pool as output. It is useful to think of Operations as falling into four functional categories (Fig. 1A): source, transformation, composition, and state Operations. *Source Operations* create the initial Pools from which all downstream Pools are constructed. The Pools they create may represent user-specified sequences loaded directly or from FASTA and VCF files, DNA sequences generated using position weight matrices (PWMs) or IUPAC motifs, random k-mers, barcodes, and so on. *Transformation Operations* modify sequence content, e.g., generating sequence variants through point mutations, insertions, deletions, recombination, or shuffling. ORF-aware Operations are provided for transformations that operate on open reading frames. *Composition Operations* combine sequences from multiple parent Pools, either by concatenating parent sequences end-to-end (the `join` Operation) or by merging multiple Pools into one Pool (the `stack` Operation). *State Operations* affect which sequences are in a Pool and how they are ordered, and include methods to select, re-order, filter, and replicate sequences. Sequences can be filtered based on sequence length, GC content, homopolymer length, sequence complexity, restriction enzyme recognition sites, or other user-defined criteria.

**Fig. 1.**
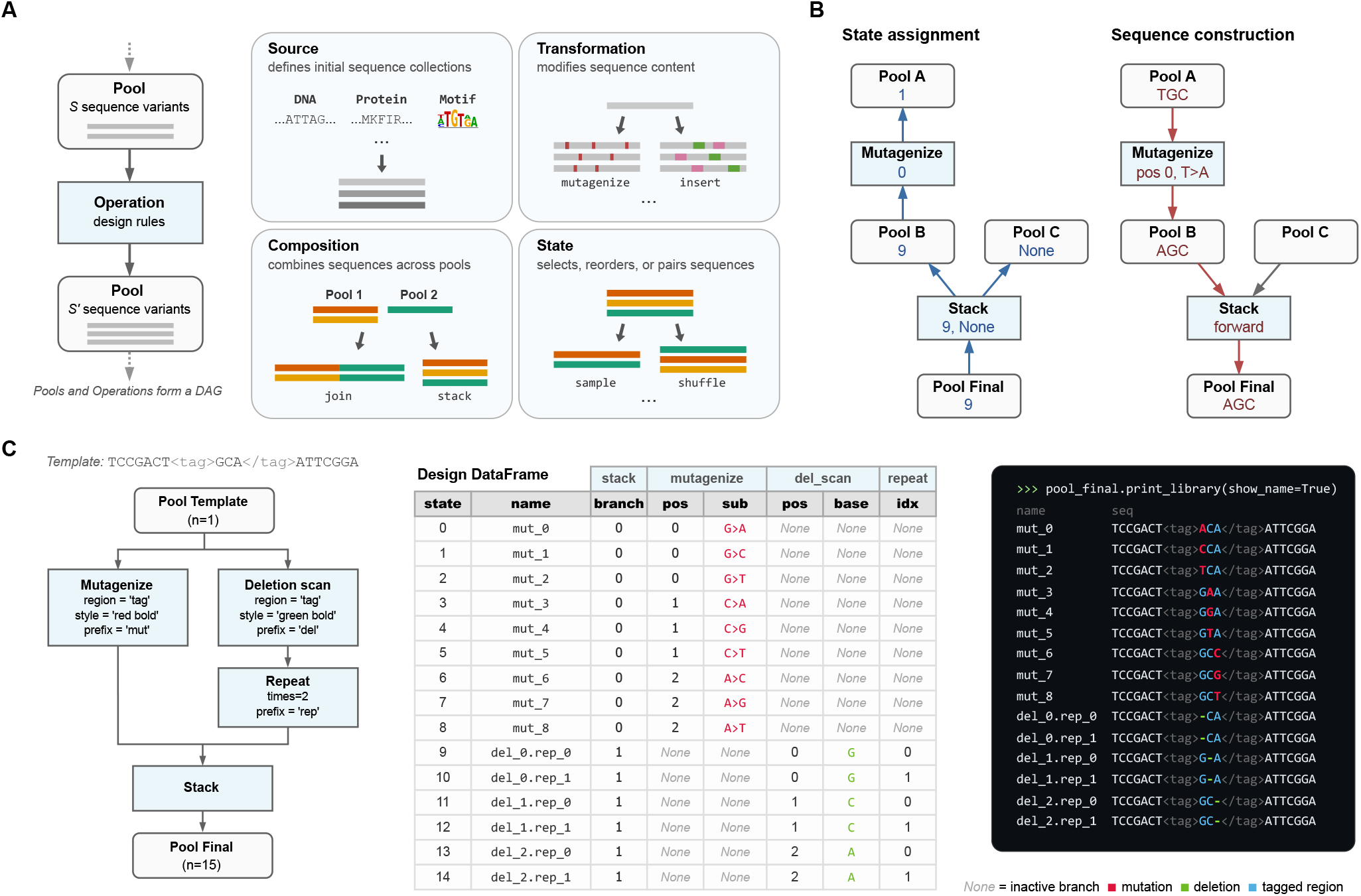
PoolParty builds libraries from Pools and Operations. **(A)** Pools represent collections of sequences, and Operations generate new Pools based on their inputs (left). Users chain Operations into a directed acyclic graph (DAG) that specifies the desired library. Operations roughly fall into four functional categories: source, transformation, composition, and state (right). **(B)** Sequence generation proceeds in two steps: state assignment and sequence construction. For each desired sequence, PoolParty passes to the root Pool a number (called the *state*) that specifies which sequence to generate. During state assignment (left), this state is propagated backward through the DAG, resolving the internal state of every Operation along the way. During sequence construction (right), each Operation generates and combines sequence fragments according to its assigned state, producing the final sequence at the root Pool. When multiple Pools are combined by `stack`, only one parent is active for a given state; inactive branches receive no state (`None`) and are shown as gray arrows. **(C)** Example workflow. The template sequence contains a user-specified region (called “tag”) that is targeted for both mutagenesis and deletion scanning. Left: the DAG specifying the sequence library. Center: the design card DataFrame returned by the DAG. Right: styled sequence output, with colors indicating mutations (red), deletions (green), and the tag region (blue).

### Sequence generation

To generate a sequence library, PoolParty iterates through a series of *states*. Each state is an integer that, together with a random seed, uniquely determines a sequence. The generation of each individual sequence proceeds in two steps: state assignment and sequence construction (Fig. 1B; Supplemental Information, Fig. S1). This process begins when the corresponding state is passed to the root Pool. State assignment involves a *backward pass* through the DAG, from the root Pool toward the source Pools. Each Pool passes the state it receives to the Operation that created it. Each Operation then decomposes this state into an *internal state* (which it keeps) and zero or more *component states*, each of which is passed to a distinct input Pool of that Operation. Sequence construction then occurs during a *forward pass* through the DAG, from the initial Pools toward the root Pool: each Operation constructs a sequence based on its internal state and the sequences returned by its input Pools, then passes this sequence to the Pool it creates. The final constructed sequence is then received from the root Pool and recorded.

### Operation modes

The specific way that each Operation interprets its internal state is determined by its *mode*.

- In *sequential mode*, the Operation exhaustively enumerates a finite set of sequence designs. For example, when `mutagenize` is instructed to generate single-nucleotide mutations in sequential mode, it yields one variant for each possible character change at each target position. Its internal state specifies which mutation to apply at which position.
- In *random mode*, the Operation samples sequence designs stochastically. For example, when `mutagenize` operates in random mode, it randomly selects which positions to mutate and which mutations to introduce. Its internal state, combined with a master seed, determines the specific random draws, ensuring reproducibility.
- In *fixed mode*, there is no internal state. Rather, output sequences are uniquely determined by the input sequences. One example is `join`, which concatenates its input sequences without modification.

### StateTracker

The way each Operation decomposes the state it receives during the backward pass is designed to reverse the way it constructs sequences during the forward pass. For most Operations, the set of possible output states is the Cartesian product of the possible internal states and the possible states of each input Pool. In such cases, the Operation uses mixed-radix decomposition to split the received state into its component states. The `stack` Operation is different: its output states are the disjoint union of its input Pool states. The `stack` Operation therefore uses the state it receives to select which input Pool to use, and to instruct this Pool (but not the other Pools) which sequence to return. Keeping track of these rules across a complex DAG requires specialized bookkeeping. We therefore developed a back-end package called StateTracker that automates state composition and decomposition during forward and backward passes through the DAG.

### Sequence regions

One may want to perform different Operations on different parts of a sequence. To facilitate this, PoolParty lets users mark regions of interest within sequences using XML-style tags. PoolParty supports both opening/closing tag pairs (e.g., `AAA<cre>CCCGGG</cre>TTT`) and self-closing tags (e.g., `ACGT<ins/>ACGT`). Operations can then be instructed to act only on a specified region, as illustrated in Fig. 1C. Tags persist through the DAG and remain valid even when upstream Operations change the content of a region. Multiple Operations can thus target modifications to the same region in series.

### Sequence metadata

It is often useful to record how each sequence within a library was constructed. PoolParty provides three complementary ways to do this (Fig. 1C).

- *Design cards* record the specific changes applied to each sequence by each Operation. Users specify which Operations and Operation-specific variables to track, including user-defined sequence scores computed with the `score` Operation. As each sequence is generated, the root Pool returns it together with a design card that records the tracked variable values for that sequence. PoolParty compiles these into a DataFrame in which each row contains a generated sequence and its design card.
- *Sequence names* summarize at a glance how each sequence was constructed. Users assign a prefix to each Operation they wish to track. PoolParty then pairs each prefix with the corresponding Operation’s internal state to form a token (e.g., `mut_03`), and concatenates these tokens to form the sequence’s name. For example, `wt.mut_03.bc_01` might denote a wild-type sequence with mutation variant number 3 and barcode number 1.
- *Sequence styling* allows sequences to be annotated with colors and formatted text that reflect each one’s construction history. For example, `mutagenize` can be instructed to mark the positions it mutates in bold yellow text, making these positions stand out from the rest of the sequence. This formatting is accomplished using ANSI escape codes, so that output renders correctly in command-line terminals, Jupyter notebooks, and other environments that support basic text formatting. These styles combine in a natural manner as sequences propagate through the DAG. For example, a mutated position can be underlined even if it occurs within a sequence region that is already colored blue.

PoolParty also provides a `stats` method for summarizing a DNA Pool, including its unique and duplicate sequence counts, pair-wise Hamming distances, and homopolymer frequencies.

### Implementation and extensibility

PoolParty requires Python 3.10 or later, is designed to have minimal dependencies, and is distributed via PyPI. Documentation, tutorials, and example notebooks are available on the ReadTheDocs and GitHub project websites. PoolParty is also extensible: users can create new Operation types by subclassing the Operation base class and implementing a few required methods. Custom Operations automatically inherit support for state tracking, design card generation, and visualization, allowing researchers to add domain-specific functionality without reimplementing core infrastructure.

### SpliceAI surrogate modeling analysis

This section provides technical details for the surrogate modeling analysis presented in Results. Using the human 5’ splice site (5’ss) position weight matrix [45], we sampled 9-mers representing cryptic 5’ss sequences and scored them with MaxEntScan [46]. Sequences were binned into 100 quantiles of MaxEntScan score (hereafter, strength), and 20 sequences were drawn from each bin (2,000 total). Each 9-mer was inserted at 100 positions flanking the canonical 5’ss of SMN2 exon 7 and paired with a matched control in which the GT dinucleotide was disrupted (T*>*A at +2).

SpliceAI [44] is a genomic deep learning model for splice-site prediction in humans. We used the five-model ensemble to predict the canonical 5’ss probability using *±*5 kb of genomic context (GRCh38). For each library-control pair, we computed the difference in logit-transformed predictions at the canonical 5’ss. These per-sequence values were averaged within each cell of the resulting 100 *×* 100 position-by-strength grid.

We fit generalized additive models (GAMs) to the cell means using pyGAM [47], with smooth terms for position, strength, and their interaction. Models were weighted by the inverse variance of each cell mean. Exonic and intronic regions were modeled separately. Model performance was evaluated by checkerboard cross-validation of the grid.

### Use of large language models

The authors used large language models (primarily Claude Opus 4.5 and 4.6, Anthropic) to assist with code development, debugging, and documentation writing. The manuscript was written entirely by the authors, with LLM assistance only at the copy-editing stage. The figures were also prepared by the authors; LLMs were used only to help write the scripts that generate the plots. The authors affirm that they are fully responsible for the content of the manuscript, the codebase, and the documentation.

## Results

### Example 1: DMS library for protein GB1

As a first example, we use PoolParty to reconstruct and extend the library used in a DMS experiment on protein GB1 by Olson et al. [17]. Their library contained the wild-type GB1 coding sequence, all single-amino-acid substitutions (excluding the start codon), and all possible pairs of amino acid substitutions. This library is substantially larger than most DMS libraries, containing over half a million variants, and has become a benchmark dataset in the quantitative modeling of protein sequence-function relationships [48–51]. To the Olson et al. library we added random higher-order amino acid substitutions as well as 100 replicates of the wild-type sequence.

Fig. 2A illustrates the DAG used to construct the resulting library; the corresponding Python code is shown in Fig. 2B. First, we use the `from_seq` Operation to define a Pool named `orf_pool` that represents the wild-type GB1 coding sequence. This Pool is then passed to four separate Operations that define the four library components. The first is a `repeat` Operation that produces 100 copies of the wild-type sequence. The second and third are `mutagenize_orf` Operations running in sequential mode: one generates all 1,045 single-amino-acid substitutions, and the other generates all 536,085 pairwise amino acid substitutions. The fourth is a `mutagenize_orf` Operation running in random mode, which mutates each amino acid position with 10% probability to generate 10,000 higher-order variants. We note that `mutagenize_orf` supports multiple codon mutation types; this example uses `missense_only_first`, which selects the codon most frequently used in the human genome for each target amino acid. These four Pools are then merged by `stack` into the root Pool, yielding a library of 547,230 sequences in total. The design card for each variant (Fig. 2C) reports which component it belongs to, the specific mutations it contains, and the replicate index (for wild-type copies). Users can also inspect the DAG structure by calling the `print_dag` method of the root Pool (Fig. 2D).

**Fig. 2.**
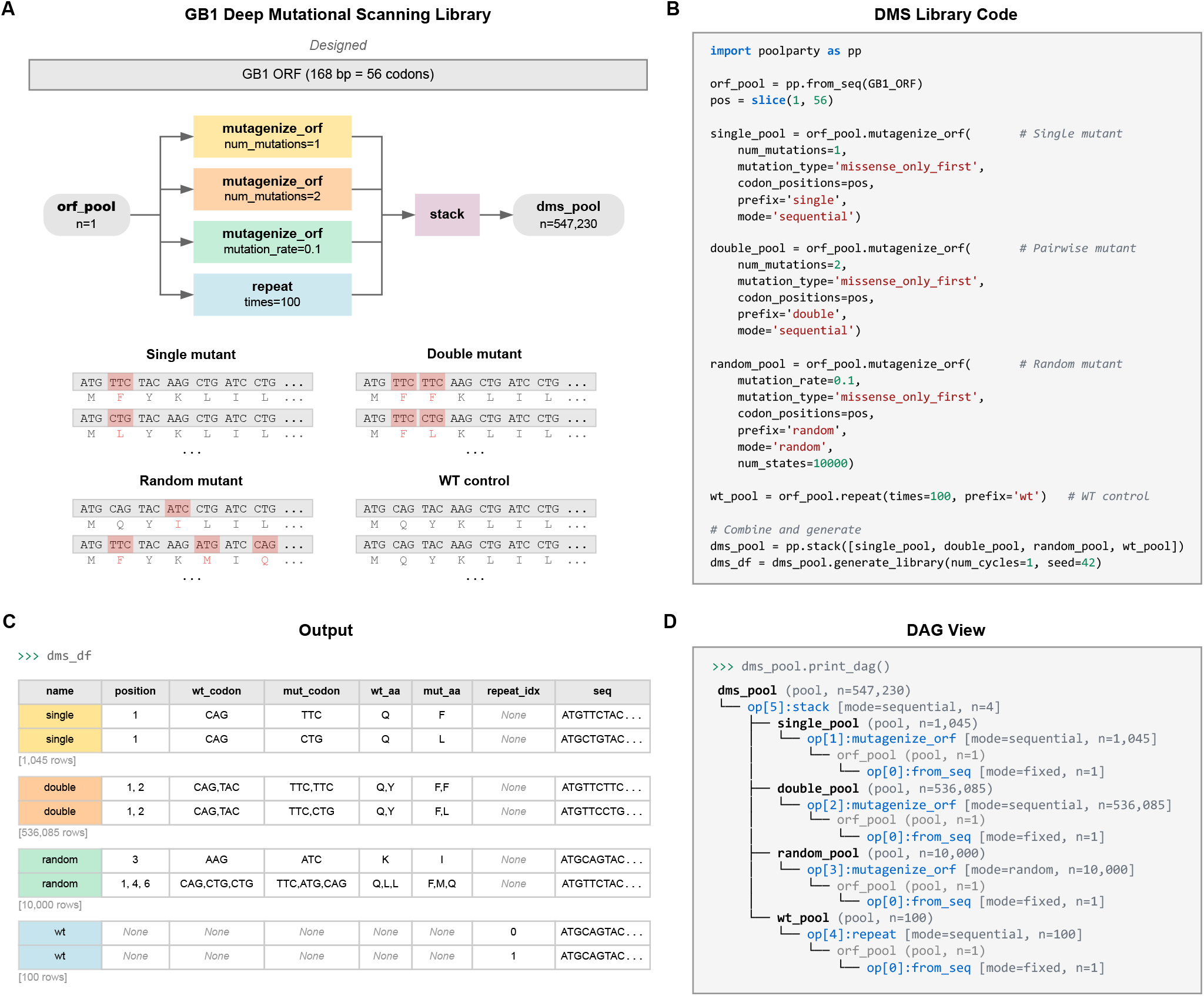
A deep mutational scanning (DMS) library for protein GB1. **(A)** DAG and representative sequences for each of the four library components: replicates of the wild-type sequence, all single-amino-acid substitutions, all pairwise amino acid substitutions, and random higher-order mutants. Mutated codons are highlighted with amino acid translations shown beneath. **(B)** Python code implementing the DAG. **(C)** Design cards reporting the library component, mutation positions, and amino acid changes for representative variants. **(D)** DAG structure displayed by `print_dag()`.

### Example 2: MPRA library for regulatory grammar

MPRA libraries used to study regulatory grammar often vary the positions and orientations of transcription factor binding sites (TF-BSs) [3, 5, 52–56]. For example, Georgakopoulos-Soares et al.[56] studied the impact of order and orientation on gene expression by testing over 200,000 sequences containing all pairwise and triplet arrangements of 18 TFBSs. Designing such libraries requires substantially more complex combinatorics than standard DMS libraries do. PoolParty handles this complexity without making additional demands on the user.

As a second example, we designed an MPRA library representing various arrangements of binding sites for three liver-enriched transcription factors—HNF4A, PPARA, and XBP1 (Fig. 3A,B). Following the oligo layout of Melnikov et al. [3], we start by creating a Pool representing a template sequence that contains a putatively inert 100 bp cis-regulatory element (CRE) region (`<cre>`) and a barcode region (`<bc>`). Next, we create a Pool representing the binding site for HNF4A. This Pool is then passed to the `flip` Operation, which yields both orientations of the site (forward and reverse complement). Pools for PPARA and XBP1 are created similarly. The template Pool and three TFBS Pools are then passed to the `insertion_multiscan` Operation, which samples 1,000 random placements of the TFBSs within the `<cre>` region without overlap. Because `flip` runs in sequential mode, all 2^3^ = 8 orientation combinations are generated for each placement, yielding 8,000 unique CRE variants. Each sequence is then replicated three times using the `repeat` Operation. Finally, the `get_barcodes` Operation is used to generate unique barcode sequences that are inserted into the `<bc>` region of the template. The result is a library of 24,000 sequences (Fig. 3A).

**Fig. 3.**
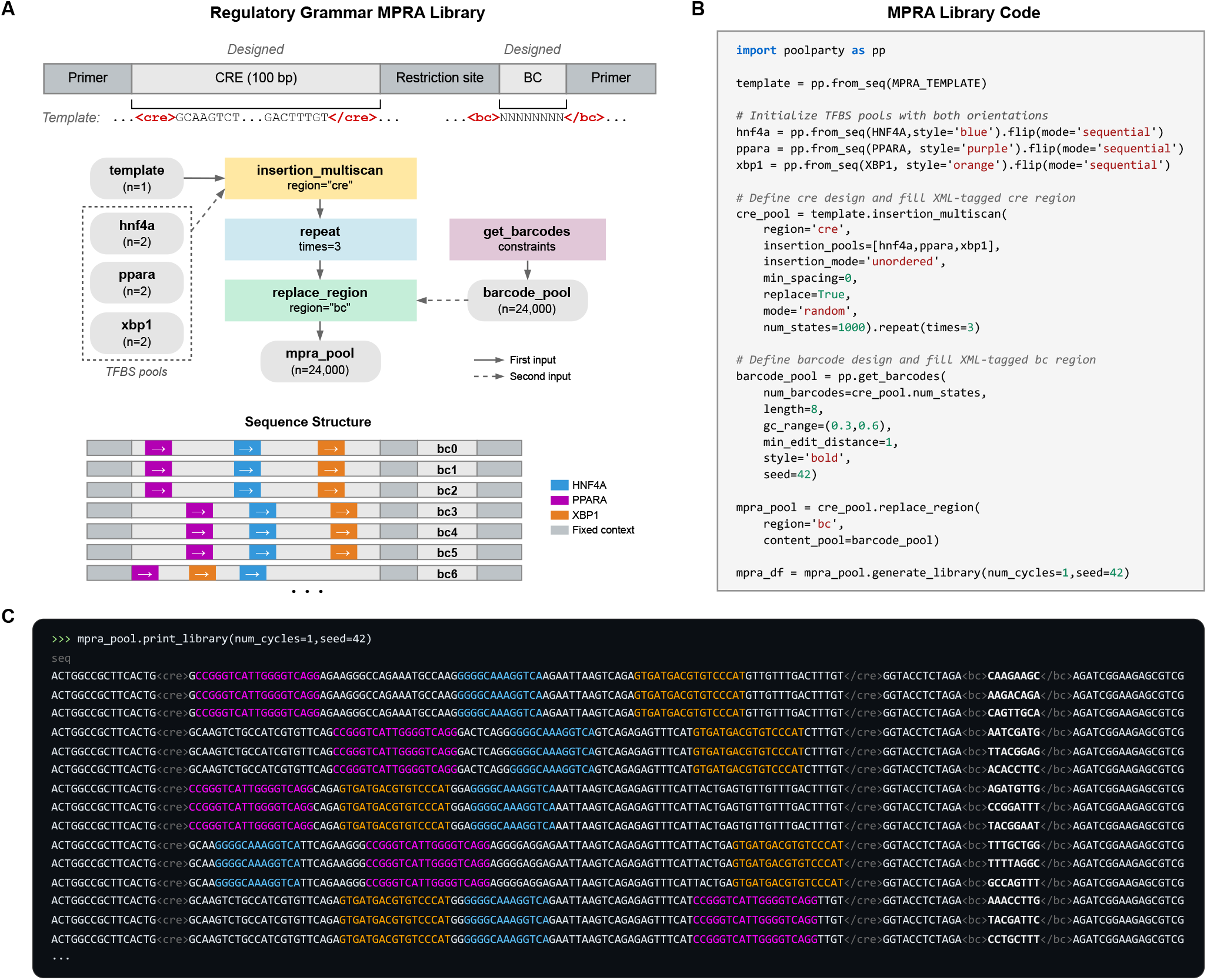
A massively parallel reporter assay (MPRA) library for probing regulatory grammar. **(A)** Library schematic showing TFBS placements across a 100-bp CRE in representative variants, each linked to three different barcodes. **(B)** Python code for library construction. TFBS Pools are initialized with forward sequences and augmented with the `flip` Operation to include both orientations. **(C)** Output of `print_library`. TFBSs are colored by identity (HNF4A blue, PPARA purple, XBP1 orange); barcodes are shown in bold.

The text styling capabilities of PoolParty help users verify the architecture of the resulting sequences. After the template Pool is created, the `stylize` Operation is used to shade the `<cre>` region gray. Each TFBS Pool is also assigned a distinct color when it is created (HNF4A blue, PPARA purple, XBP1 orange); this is specified by the `style` parameter passed to `from_seq`. Barcodes are similarly set to render in bold text (Fig. 3B). These styles compose as sequences pass through the DAG, so that when `print_library` is called (Fig. 3C), the position, order, and orientation of each binding site are immediately visible.

### Example 3: Surrogate modeling of SpliceAI predictions

Designed sequence libraries are also used to study genomic AI models [31–33, 35]. First, an in silico experiment is carried out in which the AI model scores all sequences in a library. The results are then analyzed to characterize model behavior. One useful way of analyzing these synthetic data is *surrogate modeling*, in which a mathematically simple model—one whose structure and parameters are directly interpretable—is fit to the predictions.

In addition to streamlining the design of sequence libraries, PoolParty facilitates in silico experiments on genomic AI models in two ways. First, because sequence design is separated from sequence generation, new sequences can be generated on demand as the experiment proceeds. Second, the design card that PoolParty returns with each sequence provides metadata that can be used directly as covariates in downstream analyses, including surrogate modeling.

As a third example, we carried out a surrogate modeling study of SpliceAI, a deep learning model of mRNA splicing. We asked how the insertion of a cryptic 5’ss near an existing (canonical) 5’ss affects the model’s prediction for canonical site usage. Using the 5’ss of SMN2 exon 7 as the canonical site, we used PoolParty to construct a library in which cryptic 5’ss 9-mers of varying strength (MaxEntScan score [46]) were inserted at varying positions on either side of the canonical 5’ss (see Implementation). The specific values for cryptic site strength and position were recorded in the design cards (Fig. 4A).

**Fig. 4.**
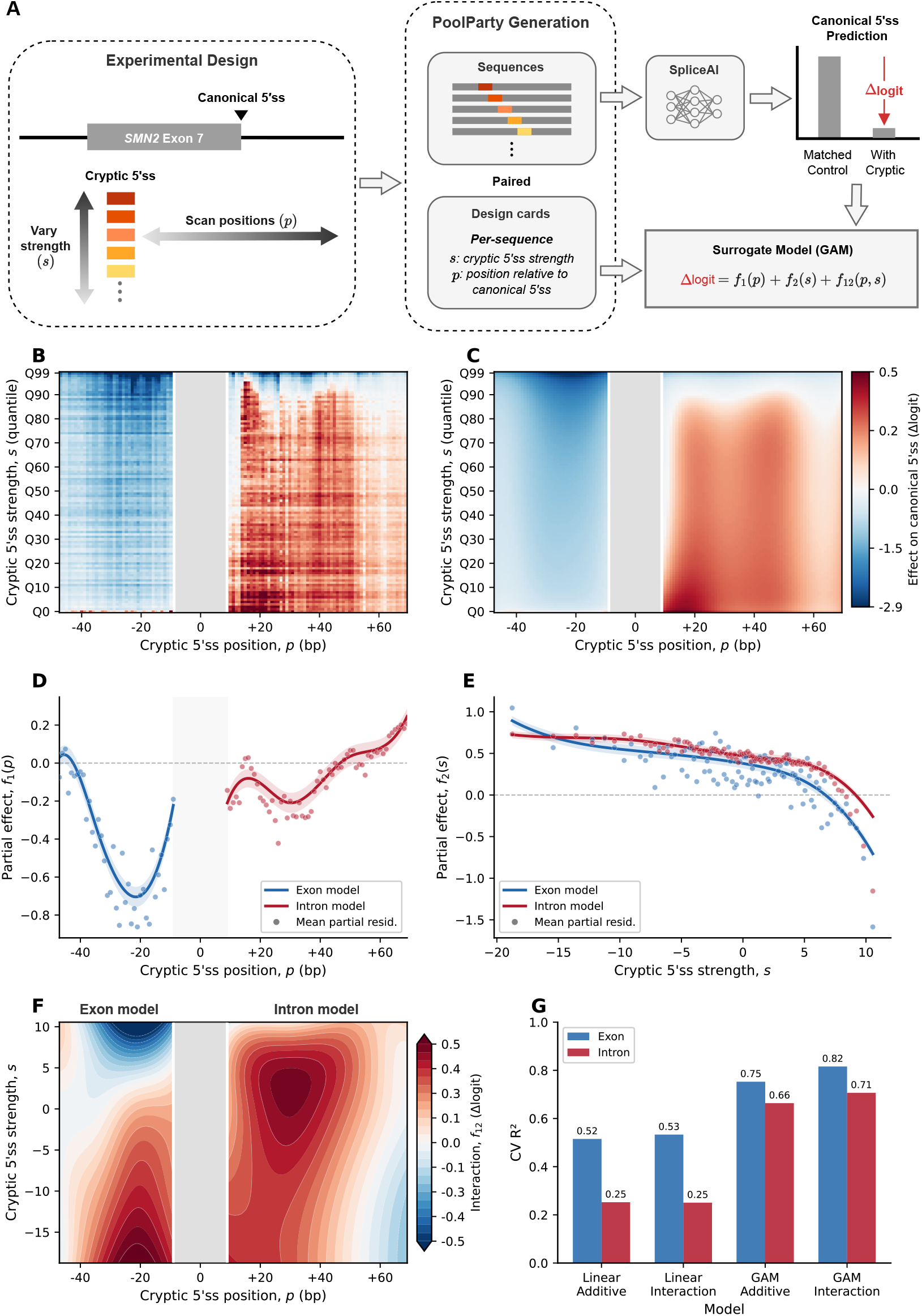
Surrogate modeling of SpliceAI predictions. **(A)** Experimental design. Cryptic 5’ss 9-mers of varying MaxEntScan strength (*s*) were inserted at varying positions (*p*) flanking the canonical 5’ss of SMN2 exon 7. Each variant was paired with a control in which the splice-site GT dinucleotide is disrupted (GT*→*GA). PoolParty records the values of *s* and *p* in the design cards. The effect of each insertion on SpliceAI predictions at the canonical 5’ss is quantified as Δlogit and modeled as a function of *s* and *p* using a generalized additive model (GAM). **(B)** Mean Δlogit across the strength *×* position grid (100*×*100; each cell averages over 20 distinct 9-mers). The gray band marks the canonical 5’ss region. **(C)** GAM predictions (including the interaction term), using the same color scale as in B. **(D)** Partial dependence of the position term *f*_1_(*p*), shown separately for exon (blue) and intron (red) models. Shaded regions indicate 95% confidence intervals; points denote mean partial residuals. **(E)** Partial dependence of the strength term *f*_2_(*s*), with conventions as in D. **(F)** Interaction surface *f*_12_(*p, s*) for exon and intron models. **(G)** Checkerboard cross-validation *R*^2^ for linear and GAM models, with and without interaction terms.

Each variant was paired with a matched control in which the cryptic site was disrupted, and the effect of each insertion was quantified as the difference in logit-transformed SpliceAI-predicted probabilities at the canonical site (Δlogit). Generalized additive models (GAMs) were used to separately model the effects of cryptic site insertion at exonic and intronic positions. Each of these two models has three terms, corresponding to the effects of position alone, strength alone, and interactions between these covariates (Fig. 4A).

The GAMs closely reproduced the observed data (Fig. 4B,C). Cryptic sites on the exonic side suppressed canonical site usage more strongly than those on the intronic side (Fig. 4D), with the effect peaking approximately 20 bp upstream of the canonical 5’ss. This is consistent with the observation that cryptic 5’ss are preferentially depleted near annotated ones on the exonic side [57]. Stronger cryptic sites drove greater suppression in both models (Fig. 4E). The effects of position and strength exhibited substantial interactions (Fig. 4F). Through a comparison to analogous linear models, we observed that the non-linear terms and interaction terms in the GAMs substantially improved predictive performance, with held-out *R*^2^ values of 0.82 and 0.71 for exonic and intronic insertions, respectively (Fig. 4G).

## Discussion

We have presented PoolParty, a Python package for designing oligonucleotide libraries. PoolParty facilitates this task by representing libraries as directed acyclic graphs (DAGs) that users can construct using a simple API. We demonstrated PoolParty through three use cases: a DMS library for protein GB1; an MPRA library probing transcriptional regulatory grammar; and a surrogate modeling analysis of the effects of cryptic 5’ss on SpliceAI predictions. These examples illustrate the flexibility of PoolParty and its applicability to both experimental and computational studies. Runtime and peak memory benchmarks for the three examples are provided in Supplemental Information (Fig. S2 and Table S1).

PoolParty was inspired by deep learning frameworks like PyTorch and TensorFlow, in which a high-level API is used to construct a DAG representing the desired neural network. The DAG then enables computation of model gradients via backpropagation. Similarly, in PoolParty, a backward pass through the DAG is used to decompose a state specific to each requested sequence into the specific Operations needed to construct that sequence during a subsequent forward pass through the DAG. This state-decomposition mechanism is a core innovation of PoolParty; in particular, it makes sequence generation random-access and allows users to test complex oligo pool designs without having to generate full sequence libraries.

We also highlighted the design card that PoolParty provides with each sequence it generates. Among other things, design cards facilitate using the parameters governing each sequence’s construction as covariates in downstream analyses. Design cards are thus a bookkeeping device that allows users to avoid reverse-engineering sequence architecture from raw sequence when investigating how this architecture affects the output of a sequence-to-function model. For example, in the SpliceAI analysis of Fig. 4, the cryptic splice-site strength and position recorded in each design card provided the covariates used for surrogate modeling. We expect design cards to be increasingly useful as in silico experiments on genomic AI models become more common.

PoolParty complements existing library-design tools such as VaLiAnT [40], MPRAnator [41], and tangermeme [43] by providing a single framework for DMS libraries, MPRA libraries, and library designs that combine elements of both. Its DAG-based structuring of oligo pools lets users freely compose arbitrary sequence generation and transformation operations. Because the DAG encodes the full construction logic, structured design-card metadata are generated automatically as a byproduct. Additional comparisons between PoolParty and existing tools are provided in Supplemental Information, Figs. S3 and S4 and Tables S2 and S3.

PoolParty was designed to solve a specific computational problem: streamlining the design of DNA sequence libraries for a wide range of experimental applications and in silico studies. As such, it has several limitations. First, PoolParty operates on sequences, not genomic loci or transcript models. It can accept genomic co-ordinates as input to extract sequences from FASTA files, but does not interpret transcript annotations (e.g., in GFF/GTF files) or keep track of genomic coordinates through subsequent design steps. This may make PoolParty less helpful for certain genome-editing applications (e.g., as compared to VaLiAnT). PoolParty is also not a sequence optimization tool; it does not design sequences that have optimized codon usage [58], that are subject to complex synthesis constraints [59], or that have optimal activity as assessed by machine learning models [60]. PoolParty also does not address the problem of physically constructing DNA sequence libraries, e.g., by designing mutagenic primers [61]. We expect, however, that PoolParty can in many cases be productively paired with complementary tools that do address these challenges.

## Conclusions

PoolParty streamlines the design of DMS experiments, MPRAs, and other multiplex assays of variant effect. As genomic AI models continue to advance, we expect that PoolParty will help researchers systematically probe and interpret the behavior of these models. Moreover, because PoolParty’s composable design makes it straightforward to define new Operations, the framework can readily be extended to support novel assay types and analysis strategies as they emerge.

## Availability and requirements

**Project name:** PoolParty

**Project home page:** https://github.com/jbkinney/poolparty-statetracker

**Documentation:** https://poolparty.readthedocs.io

**Archived version:** DOI 10.5281/zenodo.19445098

**Operating system(s):** Platform independent

**Programming language:** Python (*≥* 3.10)

**Other requirements:** NumPy (*≥* 1.20), pandas (*≥* 1.3), beartype (*≥* 0.22.9), pyfaidx (*≥* 0.8.1), typing_extensions (*≥* 4.0). The companion StateTracker package is installed automatically alongside PoolParty.

**License:** MIT License

**Any restrictions to use by non-academics:** None

## List of abbreviations

5’ss: 5’ splice site
CRE: Cis-regulatory element
DAG: Directed acyclic graph
DMS: Deep mutational scanning
GAM: Generalized additive model
MAVE: Multiplex assay of variant effect
MPRA: Massively parallel reporter assay
PWM: Position weight matrix
TFBS: Transcription factor binding site

## Declarations

### Ethics approval and consent to participate

Not applicable.

### Consent for publication

Not applicable.

### Availability of data and materials

No experimental data were generated or analyzed in this study. The sequences and software used were obtained from public sources referenced in the manuscript: the GB1 coding sequence from Olson et al. [17]; the MPRA backbone, TFBS sequences, and inert background sequence from Melnikov et al. [3] and Georgakopoulos-Soares et al. [56]; the SMN2 exon 7 genomic context from GRCh38.p13 (RefSeq assembly accession GCF_000001405.39); the human 5’ splice-site position weight matrix from Wong et al. [45]; MaxEntScan from Yeo and Burge [46]; and the pretrained SpliceAI model ensemble from Jaganathan et al. [44].

### Authors’ contributions

Z.L. conceived the project. Z.L. and J.B.K. designed the project, wrote the software, performed the analyses, and wrote the manuscript. Z.L. and A.C. wrote the documentation, developed the test suite, and deployed the software. J.B.K. supervised the research. All authors read and approved the final manuscript.

## Acknowledgements

We thank Jack Desmarais, Evan Seitz, and Kai Loell for helpful discussions.

## Supplemental Information

**Additional file 1 (PDF): Supplemental Information**. This file contains Supplemental Methods, Figs. S1–S4, and Tables S1– S3.

