## Supplemental Information for "PoolParty: streamlined design of DNA sequence libraries in Python"

September 3, 2026

### Supplemental Methods

#### Benchmarking analysis

Figure S2 and Table S1 present a benchmarking study of PoolParty using the three library designs featured in Figs. 2, 3, and 4 of the main text. Benchmarking experiments were run under Ubuntu 22.04.3 LTS in Windows Subsystem for Linux 2 on an Intel Core Ultra 9 185H workstation with 31 GB RAM allocated to WSL2, using CPython 3.12.2, PoolParty 0.1.0, and StateTracker 0.1.0. Timings are single-threaded wall-clock measurements of `generate_library`, reported as mean ± SD over five replicate runs. Peak memory is the maximum resident set size reported by `getrusage`, with each data point measured in a fresh process.

#### Tool comparisons

A literature search identified 13 tools with substantial functional overlap with PoolParty; these are listed in Table S2. Table S3 compares the specific features and capabilities of the 7 tools most similar in purpose to PoolParty. Of these 7, we judged VaLiAnT [1], MPRAnator [2], and tangermeme [3] to be the closest competitors. MPRAnator is a web application that, as of this writing, is no longer accessible. Figs. S3 and S4 provide explicit comparisons of the code/inputs required to construct identical libraries using PoolParty versus VaLiAnT (Fig. S3) or PoolParty versus tangermeme (Fig. S4). We note that neither VaLiAnT nor tangermeme is capable of generating all three of the libraries featured in main-text Figs. 2, 3, and 4.

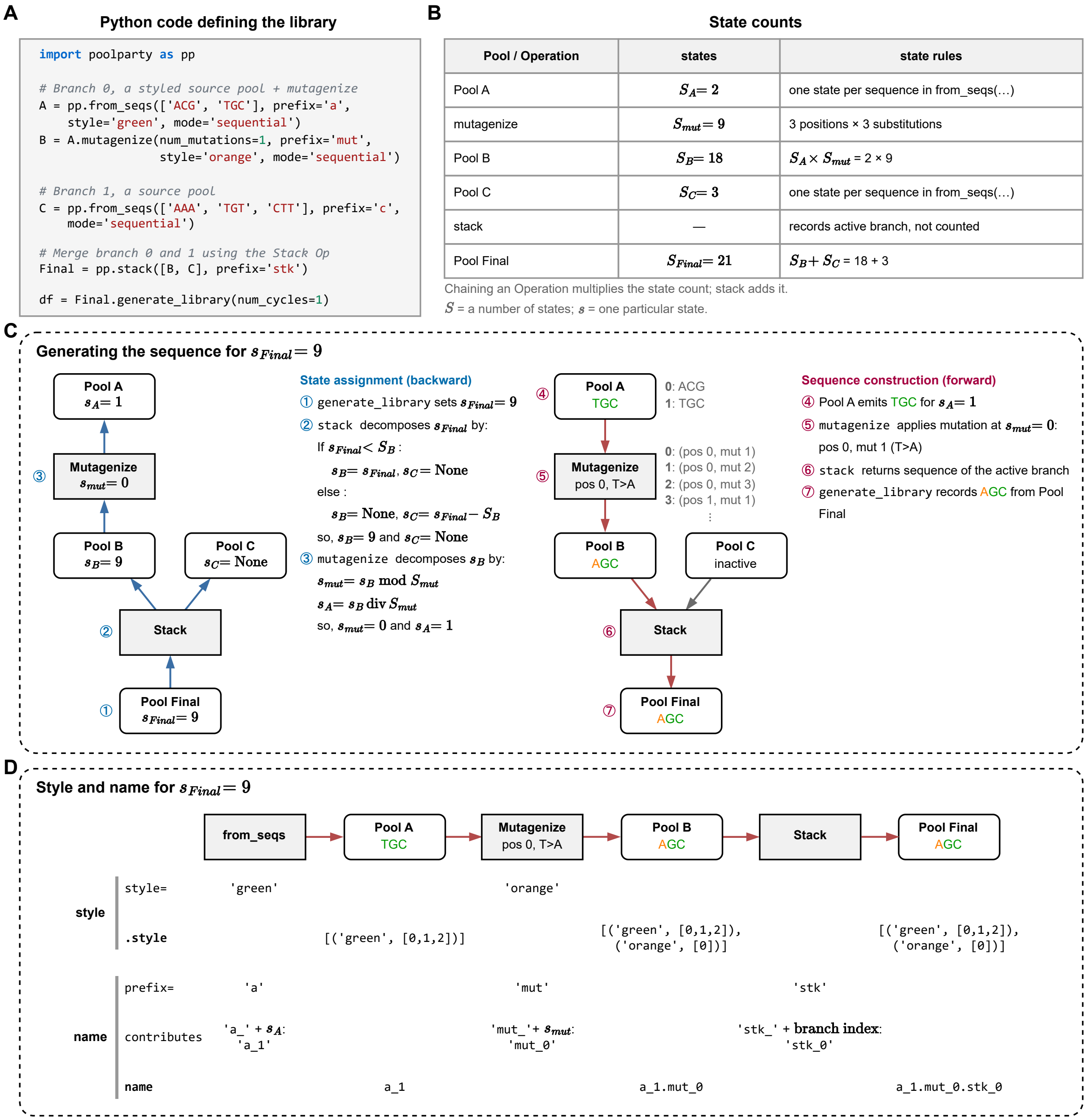

**Figure S1. A worked example of sequence generation. (A)** Python code defining the DAG in Fig. 1B of the main text. **(B)** Number of states of each Pool and Operation. **(C)** Generating the sequence for $s_{Final} = 9$. Left: state assignment (blue). Right: sequence construction (magenta). Lists beside Pool A and `mutagenize` show which sequence or mutation each state selects. Inactive branches are shown in gray. **(D)** Style and name for the same sequence. Sequences are carried through the DAG as `Seq` objects, which hold the sequence string together with its `.style`. The `style` and `prefix` arguments are supplied to Operations; the resulting `.style` and name are held by Pools. Sequence characters are colored by their `.style` entries; where entries overlap, the last one applies. The styling shown in Panel D was generated by setting `_include_inline_styles=True` in `Final.generate_library()`; this is omitted from the code snippet in panel A.

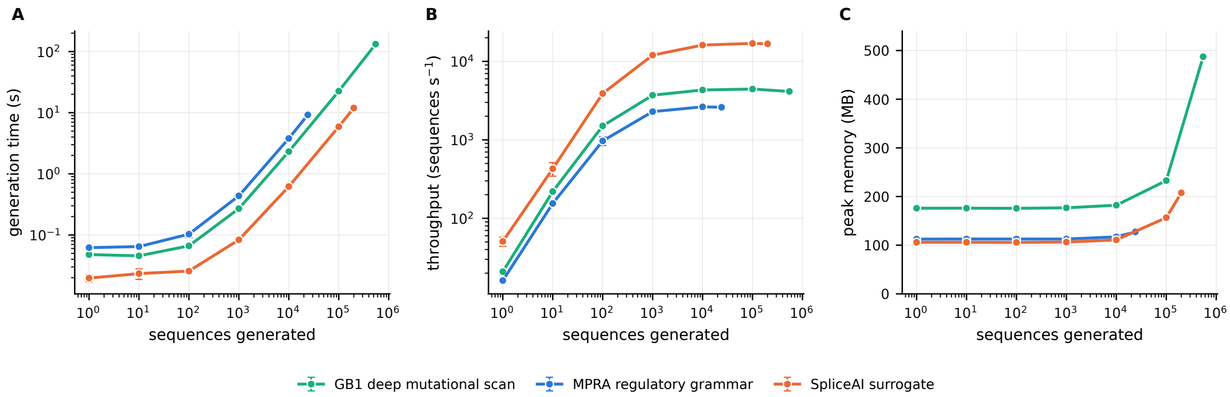

**Figure S2. Library generation scales linearly in time and memory.** **(A)** Wall-clock generation time, **(B)** throughput, and **(C)** peak resident memory against the number of sequences generated, for the three worked examples. Points are means of 5 replicate runs; error bars show $\pm 1$ SD and are smaller than the plotting symbols at most points. Below $\sim 10^3$ sequences a fixed setup cost dominates, giving the plateau in **(A)** and the rise in **(B)**; above $10^4$ sequences throughput is constant, so generation time is linear in library size. Peak memory increases linearly, by 0.52–0.66 kB per sequence above a $\sim$106 MB baseline for the MPRA and SpliceAI libraries; similar behavior is observed for the GB1 library, but with a higher memory baseline.

**A** The library: BRCA1 exon 2, three mutation types, P5/P7 adaptors

chr17:43,115,634-43,115,878 minus strand, 245 nt

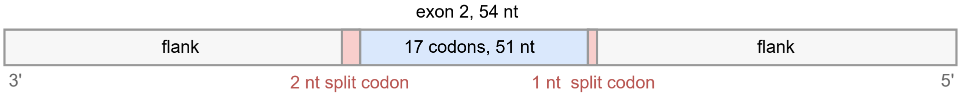

17 codons × (19 aa + 1 in-frame deletion + 1 stop) = 357 oligos

P5 20 nt + 245 nt + P7 21 nt = 286 nt (283 where a codon is deleted)

**B** PoolParty

```
import poolparty as pp

ADAPTOR_5 = "AATGATACGGCGACCACCGA"
ADAPTOR_3 = "TCGTATGCCGTCTTCTGCTTG"

# Extract region and annotate the target CDS
orf = pp.from_fasta(
    "chr17.fa", ("chr17", 43115633, 43115878, "-")
).annotate_orf(region_name="cds", extent=(100, 151), frame=1)

# Amino-acid substitutions
aa = orf.mutagenize_orf(
    region="cds", num_mutations=1,
    mutation_type="missense_only_first", mode="sequential")

# In-frame codon deletion scan
inframe = orf.deletion_scan_orf(
    deletion_codons=1, deletion_marker=None,
    region="cds", mode="sequential")

# Stop codon scan
stop = orf.insertion_scan_orf(
    "TGA", region="cds", replace=True, mode="sequential")

# Stack three pools, then add the adaptors
library = pp.join([ADAPTOR_5, pp.stack([aa, inframe, stop]),
                   ADAPTOR_3])

library_df = library.generate_library()

Inputs: 1 FASTA
Outputs: oligos, with optional design card parameters that
         produced them
```

**C** VaLiAnT

```
# Where to mutate and how
targeton.tsv
   ref_chr       chr17
   ref_strand    -
   ref_start     43115634
   ref_end       43115878
   r2_start      43115726
   r2_end        43115779
   ext_vector    "25,25"
   action_vector "(),(aa,inframe,stop),()"

# The transcript model, 51 records
ENST00000357654.9.gtf
   ...
   CDS 43115726 43115779 . - 1
   ...

# The reference genome
chr17.fa 81 MB

# Adaptors, strand, and annotation as flags
$ valiant sge targeton.tsv ../ref/chr17.fa out \
      'homo sapiens' 'GRCh38' \
      --revcomp-minus-strand \
      --adaptor-5 ... --adaptor-3 ... \
      --gff ENST00000357654.9.gtf

Inputs: 3 files
Outputs: oligos, genomic coordinates,
         consequence annotation, and a VCF
```

**Figure S3. The same BRCA1 exon 2 variant library in PoolParty and VaLiAnT.** **(A)** For each of the 17 complete codons in BRCA1 exon 2, the library contains substitutions to the other 19 amino acids, one stop-codon substitution, and one in-frame codon deletion. This gives 357 oligos. The split codons at the exon boundaries are left unchanged, and P5 and P7 adaptors are added to each sequence. **(B)** PoolParty implementation. The reference sequence is loaded from a FASTA file, and the user supplies the extent and reading frame of the 17 complete codons. Three ORF-aware Operations generate the amino-acid, stop-codon, and deletion scans. **(C)** VaLiAnT 4.0.0 implementation. VaLiAnT takes a coordinate and action table, a GTF annotation, and the reference chromosome as input. VaLiAnT gets the CDS phase from the GTF; in PoolParty, the user supplies the ORF extent and frame. The PoolParty code additionally calls `pp.set_genetic_code(valiant_codon_preference())` to match VaLiAnT's arginine codon choice; this setup line is omitted from panel B. When configured to use the same codon usage table, PoolParty and VaLiAnT gave exactly the same set of 357 sequences.

**A** The library: SMN2 exon 7, 2,000 cryptic 5'ss at 100 positions

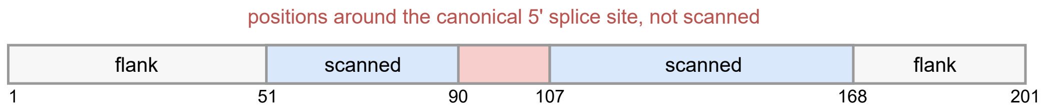

2,000 cryptic 5' splice sites × 100 positions = 200,000 sequences

```
SCAN_POSITIONS = list(range(51,90)) + list(range(107,168))
```

**B** PoolParty

```python
import poolparty as pp

# The 2,000 cryptic sites, as a pool
cryptic_pool = pp.from_seqs(
    cryptic_sites, mode="sequential")

# Replace one site at each position
library = pp.from_seq(target_region)\
    .replacement_scan(
       cryptic_pool,
       positions=SCAN_POSITIONS,
       mode="sequential")

# Final library, raw sequence strings
library_df = library.generate_library()
```

**C** tangermeme

```python
import torch
from tangermeme.ersatz import substitute
from tangermeme.utils import one_hot_encode

# The 2,000 cryptic sites, as a tensor
cryptic_ohe = torch.stack(
    [one_hot_encode(site)
     for site in cryptic_sites])

# One copy of the region per cryptic site
target_ohe = (one_hot_encode(target_region)
              .unsqueeze(0)
              .expand(len(cryptic_sites), -1, -1))

# Substitute at each position,
# Final library, one-hot encoded tensor
library_ohe = torch.stack(
    [substitute(target_ohe, cryptic_ohe, start=p)
     for p in SCAN_POSITIONS])
```

**Figure S4. The same cryptic splice-site perturbation library in PoolParty and tangermeme. (A)** The library contains 2,000 cryptic 5′ss 9-mers, each substituted at 100 positions across a 201-bp region containing the canonical 5′ss of SMN2 exon 7, giving 200,000 sequences. Positions immediately around the canonical site are excluded. This is the GT arm of the library in main-text Fig. 4; the matched control arm is built the same way. **(B)** PoolParty implementation. The 2,000 sites form a Pool, and `replacement_scan` substitutes them at the 100 positions. The generated sequences are returned as strings in a table. **(C)** tangermeme 1.4.1 implementation. The target region and cryptic sites are one-hot encoded. Substitutions are applied in one batch per position, and the final output is a tensor. After decoding this tensor, the two implementations gave the same set of 200,000 sequences.

**Table S1. Runtime and memory for the three worked examples.** Each example was generated at the size shown using a single core. Generation times are means ± SD over five replicate runs. Peak memory is the maximum resident set size and includes the fixed memory used by Python and the example DAG. DAG construction does not depend on library size and is reported separately. The SpliceAI library comprises two matched 200,000-sequence pools (cryptic GT and disrupted GA) generated separately; values are for one pool, and the full library takes twice as long.

| Example | Library size | Time (s) | Sequence/s | Memory (MB) | DAG (s) |
|---|---|---|---|---|---|
| GB1 deep mutational scan | 547,230 | $132.24 \pm 5.76$ | 4,138 | 487 | 0.22 |
| MPRA regulatory grammar | 24,000 | $9.22 \pm 0.22$ | 2,602 | 127 | 0.06 |
| SpliceAI surrogate | 200,000 | $11.93 \pm 0.27$ | 16,758 | 208 | 0.05 |

**Table S2. Tools for designing DNA sequence libraries and for related sequence-design problems.** The final two rows group related tools; their members are named and cited individually.

| Tool | Purpose / availability | Key features | Output |
|---|---|---|---|
| MPRA Design Tools [4] | Design MPRA libraries and estimate statistical power. R package (GitHub). | Barcode assignment with error-correcting sets; power analysis over effect size and sequencing depth | Barcoded constructs and statistical power estimates |
| MPRAnator [2] | Design MPRA libraries of motif arrangements and sequence variants. Web service. | Combinatorial placement of motifs; randomly sampled and combinatorial substitution sets; scrambled and reverse-complement controls | Barcoded library of sequences (FASTA) |
| Mutation Maker [5] | Design mutagenic oligos for building variant libraries in the laboratory. Web application (Docker); REST API. | Degenerate codons at user-listed sites; several substitutions combined in one reaction; de novo gene assembly with per-site variant frequencies | Primers, or gene fragments with the proportion of each variant to pool |
| Oligopool Calculator [6] | Add barcodes, primers, and spacers to a supplied variant set, and analyze the sequencing readout. Python package (PyPI); command line. | Constraint-aware barcodes, primers, and spacers; degenerate compression of a supplied variant set; off-target screening against a host genome | Synthesis-ready constructs; variant counts from sequencing reads |
| **PoolParty** | Design libraries that combine multiple variant types for laboratory assays or for probing genomic AI models. Python package (PyPI). | Libraries specified as a directed acyclic graph of Operations and Pools, allowing different mutagenesis schemes to be combined; sequences generated only on request | Library of sequences, each paired with a record of how it was constructed |
| tangermeme [3] | Generate perturbed sequence sets to probe cis-regulatory logic in deep learning models. Python package (PyPI). | Saturation mutagenesis of input sequences; insertion and removal of motifs across sets of sequences | Perturbed sequences, model predictions, or attribution scores |
| VaLiAnT [1] | Design and annotate libraries for saturation genome editing and cDNA deep mutational scanning. Command-line tool (source, Docker). | Saturation mutagenesis from genomic coordinates; several mutation types per target region; transcript-aware, including codons split across exons | Per-region libraries with metadata tables and VCF carrying variant consequence annotation |
| General-purpose toolkits *(Biopython, pydna, SeqPro)* [7–9] | Manipulate sequences, simulate cloning, prepare arrays for modeling. Python packages (PyPI). | Parsing, transformation, and file-format support | Transformed sequences |
| Sequence optimization tools *(CodonGenie, DNA Chisel, ledidi)* [10–12] | Optimize individual sequences for codon usage, synthesis constraints, or a model's objective. Python packages (PyPI); web services. | Degenerate codons covering a target residue set; hard constraints and scored objectives combined in one problem; gradient-based editing guided by a predictive model | An optimized or edited sequence, or a degenerate codon |

**Table S3. Capabilities of tools for designing DNA sequence libraries and for related sequence-design problems.** ● denotes a capability the tool provides; ◐ one provided with restrictions; ○ one the tool does not provide. Capabilities are counted only where the tool provides a dedicated operation, parameter, or mode.

| | DNA Chisel | MPRA Design Tools | MPRAnator | Mutation Maker | Oligopool Calculator | **PoolParty** | tangermeme | VaLiAnT |
|---|---|---|---|---|---|---|---|---|
| *Library Specification* | | | | | | | | |
| Library object | ○ | ◐ | ○ | ◐ | ◐ | ● | ◐ | ○ |
| Composable operations | ◐ | ○ | ◐ | ○ | ● | ● | ◐ | ○ |
| Mixed variant types in one library | ○ | ● | ● | ◐ | ○ | ● | ○ | ● |
| *Variant Generation* | | | | | | | | |
| Saturation mutagenesis | ○ | ○ | ○ | ○ | ◐ | ● | ● | ● |
| Randomly sampled variants | ● | ◐ | ● | ○ | ○ | ● | ○ | ○ |
| Pairwise and higher-order variants | ● | ○ | ● | ● | ◐ | ● | ◐ | ○ |
| Model-guided variants | ● | ○ | ○ | ○ | ○ | ◐ | ● | ○ |
| *Variant Types* | | | | | | | | |
| Codon / amino-acid substitutions | ● | ○ | ○ | ◐ | ○ | ● | ○ | ● |
| Insertions and deletions | ○ | ● | ◐ | ○ | ○ | ● | ● | ● |
| Combinatorial multi-motif placement | ○ | ○ | ● | ○ | ○ | ● | ● | ○ |
| Recombination | ○ | ○ | ○ | ○ | ○ | ● | ○ | ○ |
| Shuffling | ○ | ○ | ◐ | ○ | ○ | ● | ● | ○ |
| *Physical Construction* | | | | | | | | |
| Synthesis-constraint checking | ● | ◐ | ◐ | ○ | ● | ◐ | ○ | ◐ |
| Constraint-based optimization | ● | ◐ | ◐ | ● | ● | ○ | ○ | ○ |
| Primer design | ◐ | ○ | ○ | ● | ● | ○ | ○ | ○ |
| *Metadata and Inspection* | | | | | | | | |
| Per-sequence construction records | ● | ● | ◐ | ● | ● | ● | ○ | ● |
| Automatic naming | ○ | ◐ | ● | ● | ◐ | ● | ◐ | ● |
| Sequence styling | ○ | ○ | ◐ | ○ | ○ | ● | ○ | ○ |
| *Genomic Integration* | | | | | | | | |
| Genome coordinates | ○ | ◐ | ● | ○ | ○ | ◐ | ◐ | ● |
| Transcript / annotation aware | ○ | ○ | ○ | ○ | ○ | ○ | ○ | ● |
